# Aβ-CT affective touch: Touch pleasantness ratings for gentle stroking and deep pressure exhibit dependence on A-fibers

**DOI:** 10.1101/2022.12.06.518627

**Authors:** Laura K. Case, Nicholas Madian, Micaela V McCall, Megan L Bradson, Jaquette Liljencrantz, Benjamin Goldstein, Vince J Alasha, Marisa S Zimmerman

## Abstract

Gentle stroking of the skin is a common social touch behavior with positive affective consequences. A preference for slow versus fast stroking of hairy skin has been closely linked to the firing of unmyelinated C-tactile (CT) somatosensory afferents. Because the firing of CT afferents strongly correlates with touch pleasantness, the CT pathway has been considered a social-affective sensory pathway. Recently, ablation of the spinothalamic pathway-thought to convey all C-fiber sensations-in patients with cancer pain impaired pain, temperature, and itch, but *not* ratings of pleasant touch. This suggested integration of A and CT fiber input in the spinal cord, or A-fiber contributions to computations of touch pleasantness in the brain. However, the causal contribution of A-fibers to touch pleasantness- in humans *without* pain-remains unknown. In the current, single-blinded study we performed two types of peripheral nerve blocks in healthy adults to temporarily eliminate the contribution of A-fibers to touch perception. Our findings show that when A-fiber function is greatly diminished, the perceived intensity *and* pleasantness of both gentle stroking and deep pressure are nearly abolished. These findings demonstrate that explicit perception of the pleasantness of CT-targeted brushing and pressure both critically depend on A-fibers.

## Introduction

Gentle stroking is a prevalent affective touch behavior, whether caressing an infant, comforting a loved one, or exchanging sensual touch. While top-down effects of mood and social context strongly shape the affective nature of touch[1], there is evidence that bottom-up sensory afferents prime the affective valence of a pleasant touch experience, much as stimulation of nociceptors frequently (though not inevitably) leads to pain perception. Conventionally, myelinated Aα and Aβ afferents are understood to convey proprioceptive and touch signals to the brain, while thinly myelinated Aδ and unmyelinated C-fibers relay temperature, chemical, and pain signals[2]. However, the pleasantness of gentle stroking has been linked to a subset of C fibers called C-tactile (CT) afferents, which are maximally activated by lower-threshold tactile stimuli such as slow gentle stroking[3]. The firing of CT fibers-measured by microneurography-correlates with ratings of the pleasantness of a slow stroking stimulus[4], and CT touch activates the posterior insula[5] and increases positive affect[6]. Given their affective effects and anatomical distinction from the Aβ touch pathway, it is argued that CT fibers subserve a distinct social-affective pathway described in the “Social Touch Hypothesis” [3, 7], while A-fibers underlie more discriminative aspects of touch[7-10].

In line with the Social Touch Hypothesis, patients with hereditary reductions in C-fiber afference exhibit reduced preference for slow versus fast stroking[11]. Conversely, patients with A-fiber deafferentation show preserved CT touch pleasantness skin and insula response[5, 9], supporting its C-fiber origin. Patients with a functional loss of the *PIEZO2* ion channel subserving mechanotransduction, who exhibit accompanying severe tactile deficits, similarly remain able to detect CT-targeted slow stroking on hairy skin [12]. However, these results stem from small studies of patients with rare sensory abnormalities or disease- and may therefore be altered by abnormalities in the patients’ sensory development or compensatory brain plasticity.

Several recent findings challenge the 1:1 mapping of pleasant stroking to CT fibers. The first study demonstrates that, at least in rodents, some Aβ and CT afferents converge onto common interneurons in the spinal cord[13]. The second study, in humans, demonstrates that in patients with intractable unilateral cancer-related pain, ablation of the lamina I-spinothalamic pathway—the putative pathway for all unmyelinated afferents—largely eliminates perception of pain, temperature, and itch, but does not eliminate the pleasantness of slow stroking[14]. Finally, a study using electroencephalography (EEG) demonstrates modulation of primary somatosensory cortex by gentle stroking prior to arrival of the slower CT signal, correlated with subsequent touch pleasantness ratings. This suggests CT modulation of dorsal column (Aβ-associated) spinal projections[15]. These findings call into question the causal role of A versus CT inputs in the pleasantness of gentle stroking.

Marshall and colleagues, who performed the ablation study, propose two primary hypotheses to explain the contradictions in the CT literature[16]: 1) the alternate pathway hypothesis, in which CT fibers either join or modulate A-fibers ascending the dorsal column, and 2) the alternate percept hypothesis, in which early social touch experiences condition associations between the A- and C-fiber signals, leading to an affective response at the level of the brain[16]. In either scenario, the dorsal column pathway-historically associated with A-fiber discriminative touch-contributes to perception of affective touch pleasantness. The causal role of CT fibers in touch pleasantness in healthy humans has never been directly tested. Since A-fibers are present in the body wherever CT fibers are present, their effects are difficult to separate, except in the patient types mentioned above. However, peripheral nerve blocks present a novel opportunity to temporarily manipulate A-fiber input in healthy humans.

The affective CT theory is additionally challenged to expand by the pleasant sensation induced by deep pressure, as in massage. Our research has demonstrated the pleasant and relaxing effects of oscillating deep pressure[17], but the mechanism is not known. In humans, cutaneous anesthetic block eliminates skin sensation with little alteration in the sensation of deep pressure [18], suggesting distinct afferent pathways for deeper pressure sensation. Indeed, nerve compression blocks are reported to first block cutaneous sensation, then deep pressure, and finally deep pressure pain[19], and animal research has shown that both myelinated and unmyelinated sensory afferents in muscle can respond to muscle pressure[20-23]. Consistent with these findings, our research in humans has shown the dependence of pressure intensity sensing on Aβ afferents with a non-*Piezo2* mechanism for mechanotransduction[24]. However, the mechanism for pleasantness perception has not been studied.

Here, we conduct two types of temporary A-fiber blockade-ischemic nerve block (Study 1) and compression nerve block (Study 2)- to determine the causal contribution of A-fiber afferents in perceiving the pleasantness of two forms of affective touch. Each block technique is accompanied by specific methodological difficulties that are not shared by the other. Ischemic nerve block yields clear separation of A- and C-fiber functions[25] for a large area of skin, but cause more pain and discomfort. Nerve compression blocks affect a smaller skin surface area, but with minimal discomfort- and have been used previously to correlate specific nerve afferents with sensory percepts[26-28]. Furthermore, the latter technique has demonstrated preferential blockade of A-fibers during microneurography recordings in humans[29, 30]. Taken together, these two nerve block techniques offer complementary strengths and weaknesses that together afford a robust test of the causal contribution of A-fibers to the perception of pleasant gentle stroking and deep pressure.

## Results

### Study 1

The ischemic compression block successfully separated A- and - fiber function in the 5 participants we report data from (of the 2 participants not analyzed here, 1 reported intolerable pain and 1 lost the ability to detect heat before vibration detection was affected). By around 20 minutes, vibration detection dropped from 100% to 0 in 4/ 5 participants, and 50% in the 5^th^, while heat detection thresholds remained unaffected (<1°C change in 4 subjects, <2°C in 1). At that point in time, ratings of both intensity (previously reported in [24]) and pleasantness were nearly eliminated for both brushing (**Figure 3**) and pressure (**Figure 4**), but were largely unchanged in the control arm. Compared to baseline, nerve block reduced the pleasantness of both gentle brushing (blocked arm PRE *M* = 43.6, *SD* = 30.6, POST *M* = 3.4, *SD* = 6.5; control arm PRE *M* = 43.8, *SD* = 32.5, POST *M* = 16.0, *SD* = 14.6; *t*(4) = 3.2, *p* = 0.03, Cohen’s *d* = 0.55) and deep pressure (blocked arm PRE *M* = 19.2, *SD* = 20.0, POST *M* = 0.4, *SD* = 0.9; control arm PRE *M* = 18.6, *SD* = 21.2, POST *M* = 11.2, *SD* = 13.4, trend; *t*(4) = 2.2, *p* = 0.09, Cohen’s *d* = 0.91), as well as their intensity (gentle brushing blocked arm PRE *M* = 24.8, *SD* = 21.5, POST *M* = 3.6, *SD* = 4.6; control arm PRE *M* = 27.8, *SD* = 17.5, POST *M* = 25.2, *SD* = 19.0, trend, *t*(4) = 2.1, *p* = 0.1, Cohen’s *d* = 1.6; deep pressure blocked arm PRE *M* = 40.4, *SD* = 21.5, POST *M* = 3.2, *SD* = 7.2; control arm PRE *M* = 42.8, *SD* = 24.7, POST *M* = 30.6, *SD* = 18.4, *t*(4) = 3.3, *p* = 0.03, Cohen’s *d* = 1.7).

### Study 2

The nerve compression block successfully separated A- and C-fiber function in 17 of the 24 study participants. Seven additional subjects were dismissed from their sessions (5 reached the time limit without successful fiber separation, 1 reported intolerable pain, and 1 experienced abnormal nerve tingling prior to nerve block) and thus are not analyzed here. At about 1 hour (*M* = 52.06 min), vibration detection dropped below 50% in 17 of the 20 analyzed participants (and was maintained after affective testing in 16/17 subjects). Cold detection thresholds dropped >5°C in all 17 subjects (and were maintained after affective testing in 15/17 subjects). At that timepoint, warmth detection thresholds remained within 1°C of baseline for 12 subjects, within 2°C for 4 subjects, and within 3°C for 1 subject (and were maintained at these levels in 15/17 subjects). Participants who met all pre-established criteria for nerve fiber separation and maintained the criteria after affective testing were labelled “full responders” (*N* = 8) to the A-fiber nerve block; participants whose warmth perception rose more than 1°C or who did not maintain all criteria after affective testing were labelled “partial responders” (*N* = 9).

At the time of maximal nerve fiber separation, the intensity and pleasantness of brushing were again nearly eliminated (**Figure 3**), with significant reductions on the blocked arm relative to the control arm in both pleasantness (blocked arm PRE *M* = 31.1, *SD* = 34.0, POST *M* = 5.8, *SD* = 23.3; control arm PRE *M* = 33.8, *SD* = 3.3, POST *M* = 31.1, *SD* = 36.1, linear mixed effects model, *F*(1, 16) = 8.5, *p* = 0.01, Cohen’s *d* = 1.35) and intensity (blocked arm PRE *M* = 33.1, *SD* = 25.0, POST *M* = 5.3, *SD* = 9.3; control arm PRE *M* = 34.5, *SD* = 24.5, POST *M* = 32.8, *SD* = 23.8, *F*(1, 16) = 22.2, *p* = <0.001, Cohen’s *d* = 1.92). Similarly, the intensity and pleasantness of deep pressure were also again nearly eliminated (see **Figure 4**), with significant reductions on the blocked arm relative to the control arm in both pleasantness (blocked arm PRE *M* = 31.0, *SD* = 33.2, POST *M* = 0.5, *SD* = 25.1; control arm PRE *M* = 28.8, *SD* = 33.1, POST *M* = 25.5, *SD* = 36.8, *F*(1, 15) = 10.6, *p* = 0.005, Cohen’s *d* = 1.33) and intensity (blocked arm PRE *M* = 37.4, *SD* = 23.6, POST *M* = 10.0, *SD* = 12.9; control arm PRE *M* = 37.1, *SD* = 24.2, POST *M* = 31.8, *SD* = 22.6, *F*(1, 15) = 17.8, *p* = <0.0007, Cohen’s *d* = 1.92).

## Discussion

Recent findings[14] have questioned the exclusive dependence of the pleasantness of gentle skin stroking on C-tactile (CT) afferents, complicating the Social Touch Hypothesis [3, 7]. In the present study, we conducted two type of nerve blocks in healthy adults to selectively reduce A-but not C-fiber function and determine the effect of this loss on pleasantness ratings of CT-targeted gentle brushing. Our findings demonstrate that after loss of A-fiber sensation, the perceived intensity *and* pleasantness of gentle brushing and deep pressure are nearly abolished. In contrast, these perceptions are maintained in the control arm. These novel findings strongly suggest that A-fiber input is necessary for explicit ratings of touch pleasantness.

In Study 1, results showed a near complete loss of both intensity and pleasantness of gentle brushing and deep pressure after an ischemic nerve blockade. This method of nerve block is highly efficient at separating A- and C-fiber function, and blocks somatosensory innervation of the full lower arm. However, it causes a significant amount of discomfort and pain, leaving questions about the effect of this pain on ratings of touch pleasantness. To address this limitation, Study 2 conducted a very similar design using a nerve compression block. This block takes longer to take effect (∼1 hour) and affects a much smaller region of skin (dorsal hand near thumb and forefinger) -but does so with minimal discomfort or pain. Study 2 showed a nearly identical result: near complete loss of both intensity and pleasantness of gentle brushing and deep pressure after the nerve block. In both studies, touch pleasantness and intensity were maintained on the control arm, suggesting that results cannot be attributed to effects of the nerve block procedure on mood, or distracting effects of pain and discomfort. These techniques provide convergent evidence for the dependence of explicit touch pleasantness ratings on A-fibers.

Our results are in line with the unexpected findings of Marshall and colleagues[14, 16], who reported that ablation of the lamina I-anterolateral pathway at C1/C2 reduced perception of pain, temperature, and itch, but not the pleasantness of slow stroking[14]. The lamina I-anterolateral pathway is the putative spinal pathway for unmyelinated afferents projecting to the thalamus, as well as the spinohypothalamic and spinoparabrachial pathways. Their result suggests the sufficiency of the dorsal column pathway for explicit perception of touch pleasantness. This could be due to CT fibers joining or modulating the dorsal column pathway below the level of ablation-Marshall and colleagues’ ‘alternate pathway hypothesis’ [16].

However, our data show that C-fiber input alone is *not* sufficient for touch pleasantness perception, demonstrating a critical role of A-fibers. Our data are less clear regarding the ‘alternate percept hypothesis,’ in which early social touch experiences condition associations between A- and C-fiber signals, explaining the sufficiency of dorsal column input. Our data show that A-fibers are not just sufficient, but *necessary* for interpretation of touch pleasantness. We propose a modified ‘alternate percept hypothesis’ in which C-fibers condition responses to affective touch, but cannot be interpreted in the absence of corresponding A-fiber input.

Our results additionally demonstrate that A-fibers are critical for the interpretation of the pleasantness of deep pressure. This is less surprising, given the aforementioned association of deep pressure sensation with innervation of deeper tissues suggested by multiple animal and human studies [18-23], as well as our work demonstrating its non-*Piezo2* mechanism, contrasting with light touch sensation[24]. However, the potential contributions of CT fibers to deep pressure pleasantness are unknown.

Our findings are limited by the fact that it is not possible to fully separate A- and C-fiber function by means of nerve block. To mitigate this challenge, we have performed two methods of nerve block whose strengths and limitations complement one another. Through this approach we provide strong convergent evidence for the reliance of gentle stroking pleasantness on A-fibers. An additional limitation to our data is that participants cannot be fully blinded to the nerve block procedure; sensory changes are self-evident. However, participants were naïve to the timeline of anticipated sensory effects and were told that effects of the nerve block on many forms touch are unknown. While we demonstrate that explicit touch pleasantness ratings are highly impacted by A-fiber nerve block, it remains to be tested whether implicit measures of affective response such as facial expression are similarly impacted, confirming the dependence of the full range of CT affective effects on the contribution of A-fibers.

In sum, our convergent data from two nerve block techniques performed to block afferent A-fiber input in healthy adults suggests that in typical individuals, both A- and C-fiber afferents are important contributors to the pleasantness of CT-targeted gentle stroking and deep pressure. This study expands our understanding of the somatosensory pathways that underlie the affective and social effects of touch, and may inform future targets for noninvasive modulation of affect.

## Materials and Methods

### Study 1

#### Participants

Study 1 was approved by the NIH CNS Institutional Review Board. This study was a preliminary study and no sample size calculation was performed. Healthy controls were selected based on age and sex from participants in a broad screening protocol at NCCIH. Potential participants were scheduled for a telephone screening during which the study procedures were described, and eligibility criteria were reviewed. Participants underwent medical screening and were excluded if they had unstable medical or psychiatric conditions and any abnormalities of the skin or nerves. All participants provided informed consent and were financially compensated for their time. A total of 7 healthy volunteers participated; complete data with successful separation of A- and C-fiber nerve function was obtained and analyzed from 5 participants (2 female and 3 male, ages 21-25).

#### Methods

##### Baseline affective touch task

At baseline each participant received gentle brushing (back of the hand at a rate of 3 cm/s for 15s using a soft goat hair watercolor brush, **Figure 1a**). Participants rated each of these stimuli on two visual analog scales-one for intensity (anchors of “no sensation” (coded as 0) to “highest possible intensity” (coded as 100)) and one for pleasant/unpleasantness (anchors “extremely unpleasant” (−100) to “neutral” (0) to “extremely pleasant” (100)). Participants then received oscillating deep pressure from a commercially available hand massager (**Figure 1b)** for 20s, and rated it on the same intensity and pleasant/unpleasantness scales. Testing was conducted on the arm to be blocked and then on the control arm.

**Figure 1.**
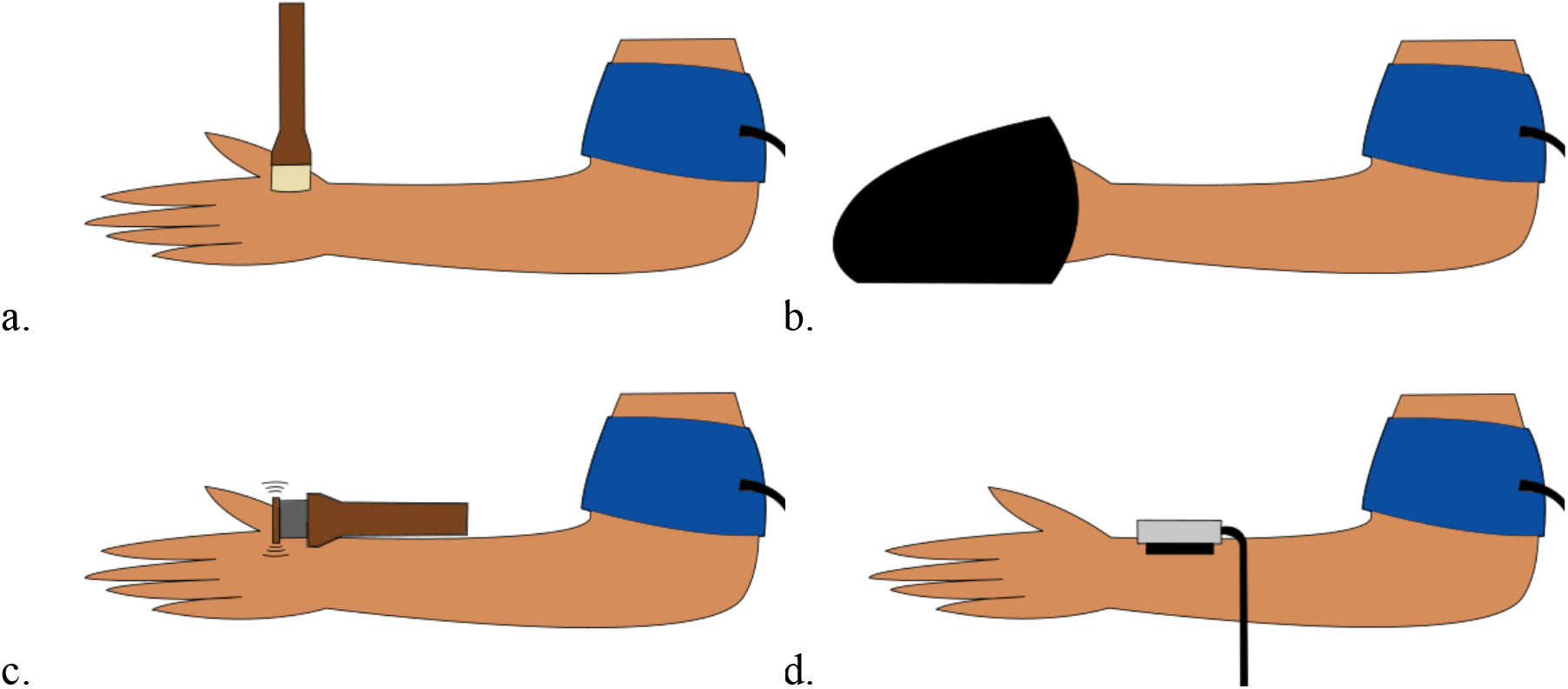
Somatosensory stimuli administered during ischemic compression nerve block. A. Gentle brushing was administered with at a rate of 3 cm/s using a soft goat hair watercolor brush. B. Deep pressure was administered using a commercially available hand massager. C. Vibration sensation was tested using a custom vibration device at 200Hz. D. Perception of warmth was tested using a Medoc thermode.

##### Nerve block placement

The participant’s arm was elevated above the head and exsanguinated for about 1 minute. Then, an automated blood pressure cuff device was wrapped around the brachium of the arm and was rapidly inflated to approximately 100mmHg above the participant’s systolic blood pressure. The arm was then rested on a pillow with the dorsal side down. Vital signs were monitored at regular intervals.

##### Nerve function monitoring

We started with four baseline rounds of testing, which included tests of several different sensory stimuli that have known associations with specific afferents. To track Aβ function we used a custom vibration device that applied 200Hz vibration for a random interval of 1-6s on a 1.3 × 4 cm region of skin on the dorsal forearm near the wrist using a custom-built probe (4.0 cm × 1.2cm × 0.7cm of balsa wood connected to a piezo-element (Piezo Systems, Inc., Cambridge, MA, USA; previously used in [31]) (**Figure 1c)**. Participants reported the onset and offset of vibration verbally over a set of three trials. To track C-fiber function we applied a Medoc thermode (Medoc, Ramat Yishay, Israel) (**Figure 1d)** over the ventral forearm at 32°C, and increased the temperature at a rate of 1°C/s until the participants indicated perception of warmth by a button press. Additional somatosensory tasks for other purposes were conducted that are not reported here. The vibration and warmth threshold tasks were repeated approximately every 2 minutes until a substantial loss of vibration detection (<50% detection) was observed.

Final affective testing: the baseline affective touch task was repeated directly after loss of vibration perception.

During all testing the participants wore noise-isolating headphones playing white noise and had a visual barrier obscuring their vision of the stimuli.

### Study 2

We initiated Study 2 to overcome limitations of Study 1, particularly the painful and aversive nature of the ischemic nerve block. Study 2 was preregistered with the Open Science Framework, doi https://osf.io/q2b68.

#### Participants

Study 2 was approved by the UC San Diego Biomedical Institutional Review Board. Given the Cohen’s *d effect sizes of* 1.3 and 1.6 in Study 1, and assuming a within-subject correlation of 0.5 and an attrition rate of 35%, a sample size of 24 was proposed to provide more than 0.8 power to detect an effect size of at least *d* = 0.8 with a two-sided α = 0.05. Healthy controls were recruited from the local university and community, and from previous studies. Potential participants were scheduled for a telephone screening during which the study procedures were described, and eligibility criteria were reviewed. Participants were included if they were 18-50 years of age, right-handed, fluent in English, and had no indication of chronic pain or current pain. Participants were excluded if they had BMI >40, unstable psychiatric conditions, current opiate use or pregnancy (urine drug screen), current lactation, history of fainting from medical procedures, allergies to latex, major medical conditions, sensory or motor abnormalities, coagulopathy or use of anti-coagulant medications, inability to communicate with investigator or rate sensations, nerve block site infection or injury, or any other medical counterindications to nerve block. All participants provided informed consent and were financially compensated for their time. A total of 24 healthy volunteers participated (7 male and 17 female; ages 20-50; self-reported ethnicity 5 White, 5 Hispanic, 6 Asian, 1 mixed; *M* = 26.8, *SD* = 7.64).

#### Methods

Participants completed a urine pregnancy test and opiate drug test.

Perception of affective touch (*Brushing Rating Task* and *Pressure Rating Task*) was tested before and after the nerve block took effect, first on the blocked arm and then on the control arm. All testing was conducted within the region of the dorsal hand affected by the compression block.

##### Brushing Rating Task

First, gentle brushing (**Figure 2a**) was administered sequentially to the blocked and control arm for 15s each, using the side of a goat hair watercolor brush (<1cm). At the end of each brushing period, participants made ratings on two visual analog scales of intensity (anchors of “no sensation” (later coded as 0) to “highest possible intensity” (100)) and pleasant/unpleasantness (anchors “extremely unpleasant” (later coded as -100) to “neutral” (0) to “extremely pleasant” (100)).

##### Pressure Rating Task

Deep pressure was administered to the blocked and control arm for 15s each using a handheld rolling massage ball (**Figure 2b**). Participants rated intensity and pleasant/unpleasantness as in the *Brushing Rating Task*.

**Figure 2.**
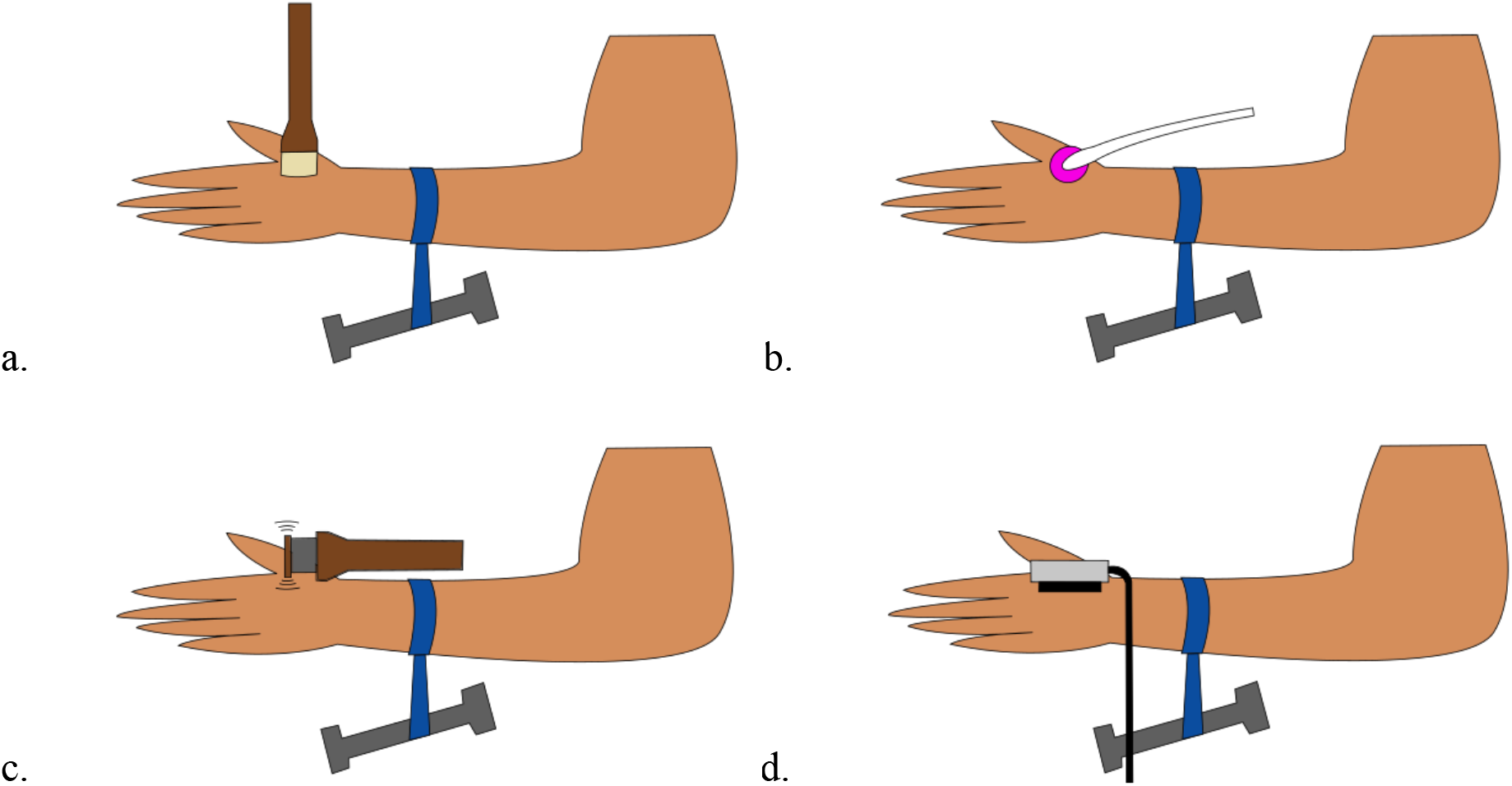
Somatosensory stimuli administered during nerve compression block. A. Gentle brushing was administered with at a rate of 3 cm/s using a soft goat hair watercolor brush. B. Deep pressure was administered using a commercially available hand massager. C. Vibration sensation was tested using a custom vibration device at 200Hz. D. Perception of cold and warmth were tested using a QST.Lab T09 thermode.

**Figure 3.**
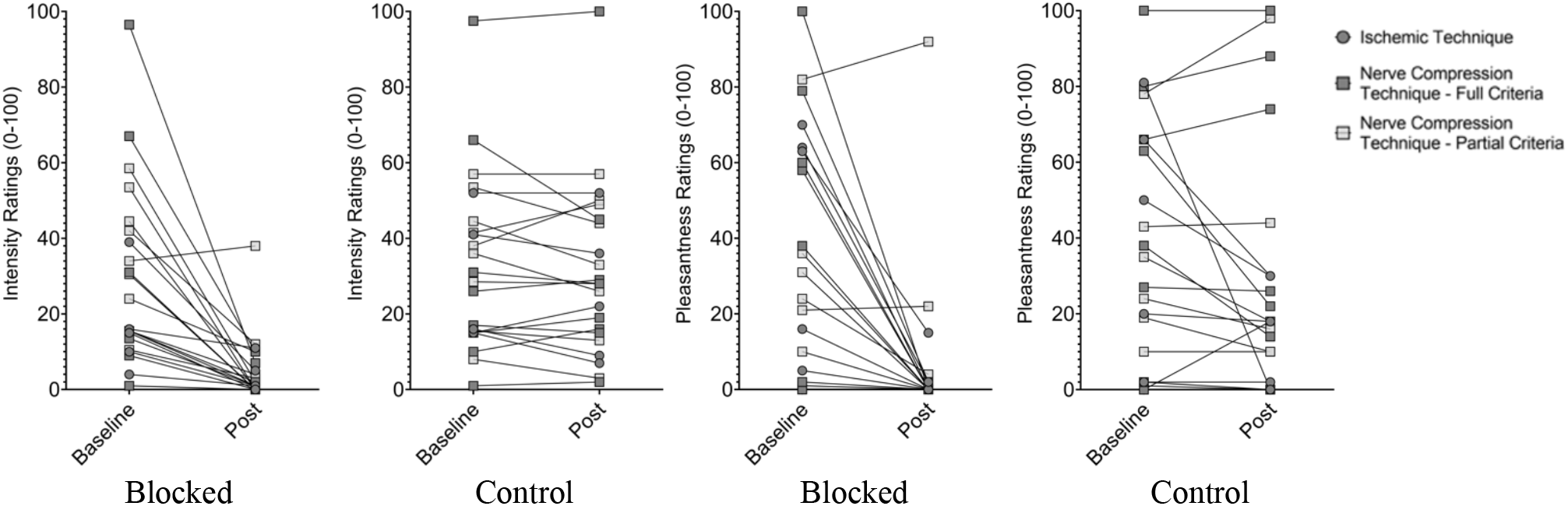
Effect of A-fiber block on intensity and pleasantness of gentle brushing. The intensity and pleasantness of slow gentle brushing on the hand or arm was rated after ischemic or compression nerve block, upon sufficient loss of A-fiber associated sensation. Participants who met all pre-established criteria for nerve fiber separation and maintained the criteria after affective testing are labelled “full responders”; participants whose warmth perception rose more than 1°C or who did not maintain all criteria directly after the brushing task are labelled “partial responders”.

**Figure 4.**
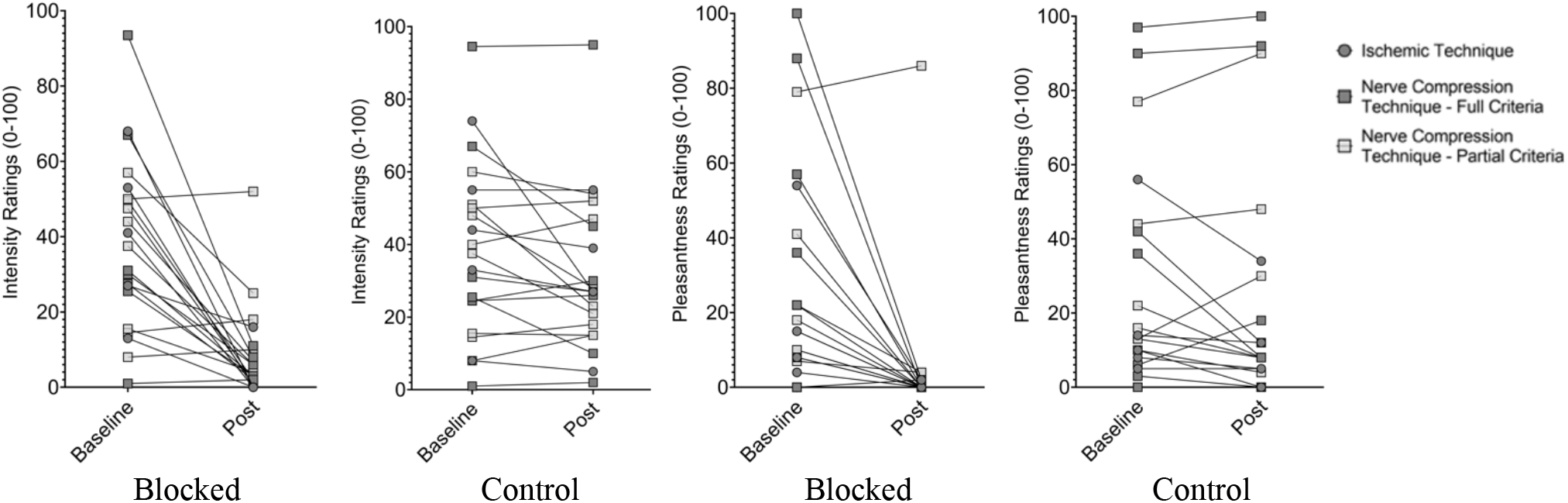
Effect of A-fiber block on intensity and pleasantness of deep pressure. The intensity and pleasantness of deep pressure was rated after ischemic or compression nerve block, upon sufficient loss of A-fiber associated sensation. Participants who met all pre-established criteria for nerve fiber separation and maintained the criteria after affective testing are labelled “full responders”; participants whose warmth perception rose more than 1°C or who did not maintain all criteria directly after the brushing task are labelled “partial responders”.

##### Baseline Nerve Function Tasks

Cold detection, vibration detection, and warmth detection were assessed at baseline, before placement of the nerve block. Each task was comprised of three trials and the mean of the three trials was taken to establish baseline sensory function. The same vibration task was used as in Study 1, but vibration was applied to the dorsal hand (**Figure 2c**). In the cold detection task, a QST.Lab T09 thermode (QST.Lab, Strasbourg, France) was placed on the dorsal hand in the area anticipated to be blocked (**Figure 2d**). The thermode started at the participant’s skin temperature and was lowered at a rate of 2°C/s until the participant indicated their perception of a cooling sensation via a response button. In the warmth detection task, the thermode was placed on the dorsal hand and increased at a rate of 2°C/s until the participant indicated their perception of a warming sensation.

The *Brushing Rating Task, Pressure Rating Task*, and *Baseline Nerve Function Tasks* were each conducted a second time to provide familiarization and comfort with the tasks prior to nerve block placement.

##### Nerve Block Placement

We initiated a nerve compression block over the left superficial radial nerve following validated procedures[26, 32, 33]: while the left hand rested in semi-prone position, a ∼1-inch cloth tourniquet was placed over the left forearm about 7cm from the wrist. A 5-lb weight was dangled from the tourniquet, similar to the weights used in some nerve compression studies[27] (see **Figure 2**). This technique often takes an hour to achieve loss of touch and cold perception[33], but does not affect major blood vessels or induce significant pain[28]. The block was released within a common safety time window of 90min for healthy research participants[26].

##### Nerve Function Monitoring

After block placement, cold and warm detection thresholds were monitored every ∼5 minutes following the same procedure as at baseline. The first two rounds of monitoring were used to establish baseline sensory nerve function. The function of Aβ fibers was monitored by the vibration task and cold threshold, with a loss of A-fiber function determined by vibration perception <50% (as in our previous study [24]), and a drop in cold threshold of >5°C. The anesthetic zone was monitored with a cotton swab, given variability in distribution of the superficial radial nerve[34], and stimulus placement was adjusted accordingly. The continued function of C-fibers was confirmed by warm thresholds maintained within 1°C of baseline[27].

##### Post-Block Affective Touch Testing

The *Brushing Ratings Task* and *Pressure Rating Task* were repeated directly after the loss of vibration and cold detection.

##### Final Nerve Function Confirmation

After the nerve block was achieved and the affective touch testing was completed, a final round of nerve function testing was conducted to confirm maintained loss of A-fiber sensation and preservation of C-fiber function.

Upon completion of all test procedures *or* upon reaching the 90min safety limit, the tourniquet was removed, and sensory function was quickly restored to baseline.

##### Data analysis

Study 1 was a preliminary study with lower power. We conducted paired t-tests to compare ratings of pleasantness and intensity before versus after nerve block. In Study 2, we conducted linear mixed effect analyses using pleasantness and intensity as dependent measures, time, arm, and their interaction as fixed effects, and participant intercept and slopes as random effects.

## Acknowledgements

We are indebted to Catherine Bushnell for her mentorship and supervision at NCCIH, and her feedback on this manuscript. We are also deeply grateful to the clinicians and staff at NCCIH for assistance in conducting Study 1. Additionally, we are grateful to Dr. Olausson for the use of his custom-built vibration devices.

The project described was partially supported by the National Institutes of Health, Grant UL1TR001442, the Intramural Research program of the National Center for Complementary and Integrative Health (NCCIH)–National Institutes of Health, and NCCIH **Grant** R00-AT009466. The content is solely the responsibility of the authors and does not necessarily represent the official views of the NIH.

## Data Availability

Data are available upon request from the corresponding author.

## References

1. Sailer, U., and Leknes, S. (2022). Meaning makes touch affective. Current Opinion in Behavioral Sciences 44, 101099.

2. Burgess, P.R., and Perl, E. (1967). Myelinated afferent fibres responding specifically to noxious stimulation of the skin. The Journal of physiology 190, 541–562.

3. Vallbo, Å., Olausson, H., and Wessberg, J. (1999). Unmyelinated afferents constitute a second system coding tactile stimuli of the human hairy skin. Journal of Neurophysiology 81, 2753–2763.

4. Löken, L.S., Wessberg, J., McGlone, F., and Olausson, H. (2009). Coding of pleasant touch by unmyelinated afferents in humans. Nature neuroscience 12, 547–548.

5. Olausson, H., Lamarre, Y., Backlund, H., Morin, C., Wallin, B., Starck, G., Ekholm, S., Strigo, I., Worsley, K., and Vallbo, Å. (2002). Unmyelinated tactile afferents signal touch and project to insular cortex. Nature neuroscience 5, 900–904.

6. Pawling, R., Trotter, P.D., McGlone, F.P., and Walker, S.C. (2017). A positive touch: C-tactile afferent targeted skin stimulation carries an appetitive motivational value. Biological Psychology 129, 186–194.

7. Morrison, I., Loken, L.S., and Olausson, H. (2010). The skin as a social organ. Exp Brain Res 204, 305–314.

8. McGlone, F., Vallbo, A.B., Olausson, H., Loken, L., and Wessberg, J. (2007). Discriminative touch and emotional touch. Can J Exp Psychol 61, 173–183.

9. Olausson, H., Cole, J., Rylander, K., McGlone, F., Lamarre, Y., Wallin, B.G., Krämer, H., Wessberg, J., Elam, M., and Bushnell, M.C. (2008). Functional role of unmyelinated tactile afferents in human hairy skin: sympathetic response and perceptual localization. Experimental Brain Research 184, 135–140.

10. Gordon, I., Voos, A.C., Bennett, R.H., Bolling, D.Z., Pelphrey, K.A., and Kaiser, M.D. (2013). Brain mechanisms for processing affective touch. Human Brain Mapping 34, 914–922.

11. Morrison, I., Löken, L.S., Minde, J., Wessberg, J., Perini, I., Nennesmo, I., and Olausson, H. (2011). Reduced C-afferent fibre density affects perceived pleasantness and empathy for touch. Brain 134, 1116–1126.

12. Chesler, A.T., Szczot, M., Bharucha-Goebel, D., Čeko, M., Donkervoort, S., Laubacher, C., Hayes, L.H., Alter, K., Zampieri, C., and Stanley, C. (2016). The role of PIEZO2 in human mechanosensation. N Engl J Med 2016, 1355–1364.

13. Abraira, V.E., Kuehn, E.D., Chirila, A.M., Springel, M.W., Toliver, A.A., Zimmerman, A.L., Orefice, L.L., Boyle, K.A., Bai, L., and Song, B.J. (2017). The cellular and synaptic architecture of the mechanosensory dorsal horn. Cell 168, 295-310. e219.

14. Marshall, A.G., Sharma, M.L., Marley, K., Olausson, H., and McGlone, F.P. (2019). Spinal signalling of C-fiber mediated pleasant touch in humans. Elife 8, e51642.

15. Schirmer, A., Lai, O., McGlone, F., Cham, C., and Lau, D. (2021). Discriminative and affective touch converge: Somatosensory cortex represents Aß input in a CT-like manner. bioRxiv.

16. Marshall, A.G., and McGlone, F.P. (2020). Affective touch: the enigmatic spinal pathway of the C-tactile afferent. Neuroscience insights 15, 2633105520925072.

17. Case, L.K., Liljencrantz, J., McCall, M.V., Bradson, M., Necaise, A., Tubbs, J., Olausson, H., Wang, B., and Bushnell, M.C. (2020). Pleasant Deep Pressure: Expanding the Social Touch Hypothesis. Neuroscience.

18. Graven-Nielsen, T., Mense, S., and Arendt-Nielsen, L. (2004). Painful and non-painful pressure sensations from human skeletal muscle. Experimental brain research 159, 273–283.

19. Kellgren, J.M. AJ (1948). On the Behaviour of Deep and Cutaneous Sensibility During Nerve Blocks. Clinical Science 7, 1–11.

20. Kaufman, M.P., Longhurst, J.C., Rybicki, K.J., Wallach, J.H., and Mitchell, J.H. (1983). Effects of static muscular contraction on impulse activity of groups III and IV afferents in cats. Journal of Applied Physiology 55, 105–112.

21. Mense, S., and Meyer, H. (1985). Different types of slowly conducting afferent units in cat skeletal muscle and tendon. The Journal of Physiology 363, 403–417.

22. Abrahams, V. (1986). Group III and IV receptors of skeletal muscle. Canadian journal of physiology and pharmacology 64, 509–514.

23. Lewin, G.R., and McMahon, S.B. (1991). Physiological properties of primary sensory neurons appropriately and inappropriately innervating skeletal muscle in adult rats. Journal of neurophysiology 66, 1218–1231.

24. Case, L.K., Liljencrantz, J., Madian, N., Necaise, A., Tubbs, J., McCall, M., Bradson, M.L., Szczot, M., Pitcher, M.H., and Ghitani, N. (2021). Innocuous pressure sensation requires A-type afferents but not functional PIEZO2 channels in humans. Nature Communications 12, 1–10.

25. Laursen, R.J., Graven-Nielsen, T., Jensen, T.S., and Arendt-Nielsen, L. (1999). The effect of differential and complete nerve block on experimental muscle pain in humans. Muscle & Nerve: Official Journal of the American Association of Electrodiagnostic Medicine 22, 1564–1570.

26. Forstenpointner, J., Binder, A., Maag, R., Granert, O., Hüllemann, P., Peller, M., Wasner, G., Wolff, S., Jansen, O., and Siebner, H.R. (2019). Neuroimaging Of Cold Allodynia Reveals A Central Disinhibition Mechanism Of Pain. Journal of pain research 12, 3055.

27. Wahren, L.K., Torebjörk, E., and Jörum, E. (1989). Central suppression of cold-induced C fibre pain by myelinated fibre input. Pain 38, 313–319.

28. Wasner, G., Schattschneider, J., Binder, A., and Baron, R. (2004). Topical menthol—a human model for cold pain by activation and sensitization of C nociceptors. Brain 127, 1159–1171.

29. Torebjörk, H., and Hallin, R. (1973). Perceptual changes accompanying controlled preferential blocking of A and C fibre responses in intact human skin nerves. Experimental brain research 16, 321–332.

30. Mackenzie, R.A., Burke, D., Skuse, N.F., and Lethlean, A.K. (1975). Fibre function and perception during cutaneous nerve block. Journal of Neurology, Neurosurgery & Psychiatry 38, 865–873.

31. Liljencrantz, J., Strigo, I., Ellingsen, D.M., Krämer, H., Lundblad, L.C., Nagi, S.S., Leknes, S., and Olausson, H. (2017). Slow brushing reduces heat pain in humans. European Journal of Pain 21, 1173–1185.

32. Ziegler, E., Magerl, W., Meyer, R., and Treede, R.-D. (1999). Secondary hyperalgesia to punctate mechanical stimuli. Brain 122, 2245–2257.

33. Nahra, H., and Plaghki, L. (2003). The effects of A-fiber pressure block on perception and neurophysiological correlates of brief non-painful and painful CO2 laser stimuli in humans. European Journal of Pain 7, 189–199.

34. Keplinger, M., Marhofer, P., Moriggl, B., Zeitlinger, M., Muehleder-Matterey, S., and Marhofer, D. (2018). Cutaneous innervation of the hand: clinical testing in volunteers shows high intra-and inter-individual variability. British Journal of Anaesthesia 120, 836–845.

